## Supplemental File for "Neural Networks for self-adjusting Mutation Rate Estimation when the Recombination Rate is unknown"

### Supplementary Note A: Feature Importance of Neural Networks

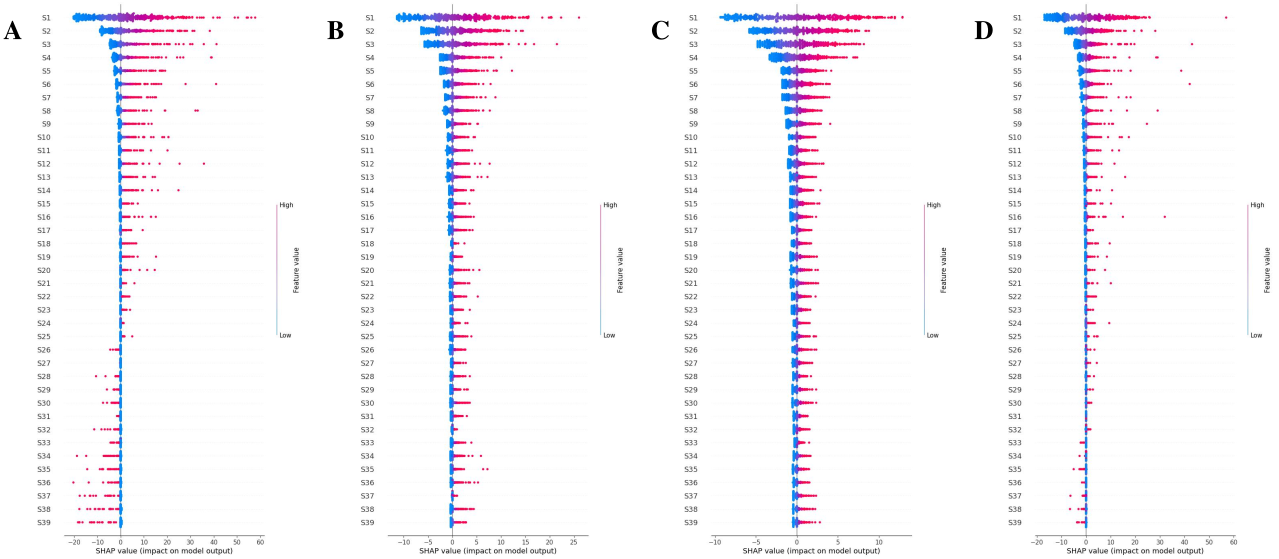


**Fig. 7. Feature importance linear neural network via SHAP.** Feature importance of the linear neural network for $\rho = 0$ (A), $\rho = 35$ (B), $\rho = 1000$ (C) and $\rho$ variable (D). Here, the calculated shap values for $\rho = 0$ refer to the linear neural network trained for $\rho = 0$ and so on. The shap values were calculated for $500$ SFS. Each dot represents the shap value for a particular prediction of one feature, i.e. one SFS entry of one of the 40 SFS. The color indicates whether the particular SFS entry had a high or low value, and the position on the x-axis shows its significance, i.e., the extent to which it increased or decreased the $\theta$ estimate.

Artificial neural networks tend to have complex architectures, which can impair the comprehension of the underlying learning process. However, it is often of interest to understand what the neural network has learned. Depending on the ML method, there are different ways to obtain knowledge about it. In our case, we determined the feature importance, i.e. a score for each feature representing its effect on the model. One tool to do so is SHAP (SHapley Additive exPlanations), a game theoretic approach to explain the output of machine learning models (29). Hereby, the shap values describe the responsibility of an SFS entry for a change in the $\theta$ estimate, the higher the absolute shap value, the greater the impact.

In Fig 7 the feature importance of the linear neural network is displayed. In general the first entry of the SFS, $S_1$, is the one with the highest impact on the estimation as its shap value is the most pronounced. For $\rho = 0$ the importance of $S_2, S_3, \ldots$ more and more decreases and increases again for the last SFS entries. For $\rho = 35$ and $\rho = 1000$, the feature importance also decreases initially, but is quasi-constant for middle and posterior $S_i$. For variable recombination rates (D), we observe a mix of the feature importance for $\rho = 0$ and $\rho = 35, 1000$. The pattern from A can still be seen, i.e. that the impact of the last SFS entries increases again and high feature values lead to a reduction of the $\theta$ estimation, but to a lesser extent. Subplots A-D in Fig 7 have in common that they show a higher importance for the first SFS entries than to the others. This is not surprising as it means that the estimator puts more weight on the mutation events happening in the more recent past, just as Fu's estimator does. Considering the weights of the linear neural network, which can be compared to the coefficients of the model-based estimators by nature, we can also extract information about the sign of the coefficient by Fig 7. For the linear neural network the weights corresponding to $S_{26}, \ldots, S_{39}$ are negative for $\rho = 0$. This can also be observed in Fig 7 A as higher values for SFS entries lead to an decreased $\theta$ estimation. Those negative weights are not surprising as half of the coefficients of the optimal model-based estimator, Fu's estimator, are negative for large enough $\theta$, see Fig S4. For $\rho = 1000$ the linear neural network has no negative weights, which can also be observed in Fig 7. This is also reasonable since Watterson's estimator has constant positive coefficients and is the optimal estimator for high recombination rates.
Fig 8 displays the feature importance for the adaptive neural network. Compared to Fig 7, no too big differences are noticeable. This shows that the adaptive neural network distributes the importance between the SFS entries more or less the same as the linear neural network, at least when trained for fixed recombination rates.
Fig. 9 shows the feature importance for the adaptive neural network trained for variable recombination rates. Shap values were calculated as before for $\rho = 0, \rho = 35, \rho = 1000$, and $\rho$ variable. By doing so, we can observe, how the adaptive network adapts to the different recombination scenarios. Again, the first entries of the SFS have the most importance, especially $S_1$, since their shap values are the most pronounced. In addition, the adaptive neural network adapts to different recombination scenarios, as we

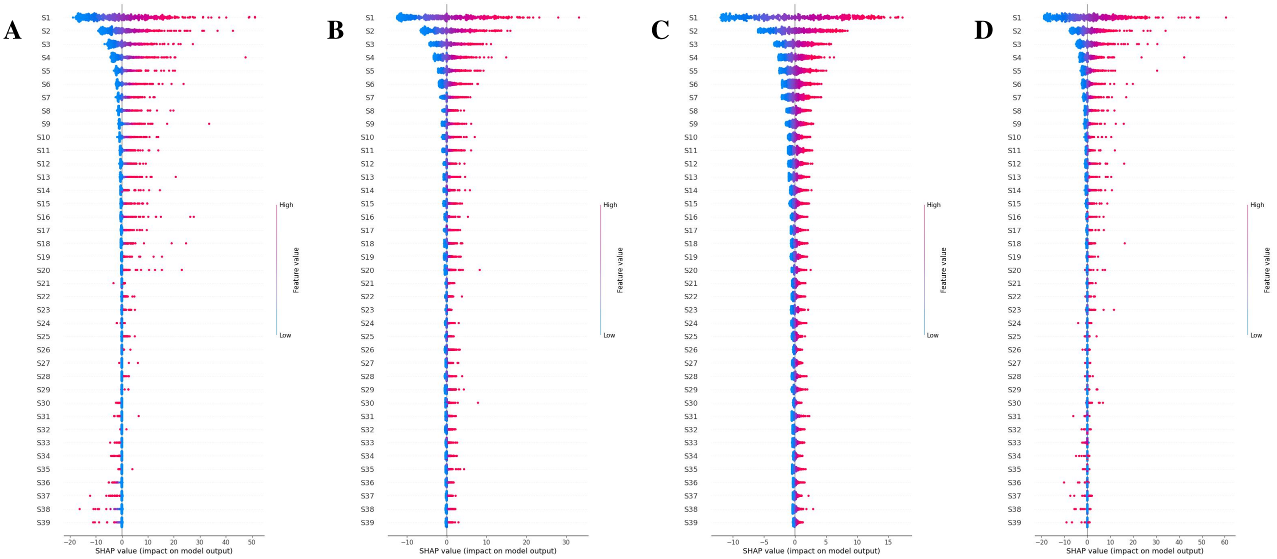


**Fig. 8. Feature importance adaptive neural network via SHAP.** Feature importance of the adaptive neural network for $\rho = 0$ (A), $\rho = 35$ (B), $\rho = 1000$ (C) and $\rho$ variable (D). Here, the calculated shap values for $\rho = 0$ refer to the adaptive neural network trained for $\rho = 0$ and so on. The shap values were calculated for $500$ SFS. Each dot represents the shap value for a particular prediction of one feature, i.e. one SFS entry of one of the 40 SFS. The color indicates whether the particular SFS entry had a high or low value, and the position on the x-axis shows its significance, i.e., the extent to which it increased or decreased the $\theta$ estimate.

can observe, for example, from a different pattern for $S_{31}$ to $S_{39}$ between subplot A and subplot B/C in Fig 9. For these features, a high value for $\rho = 0$ results in a decreased $\theta$ estimate, but for $\rho = 35$ or $\rho = 1000$ in an increased estimate.

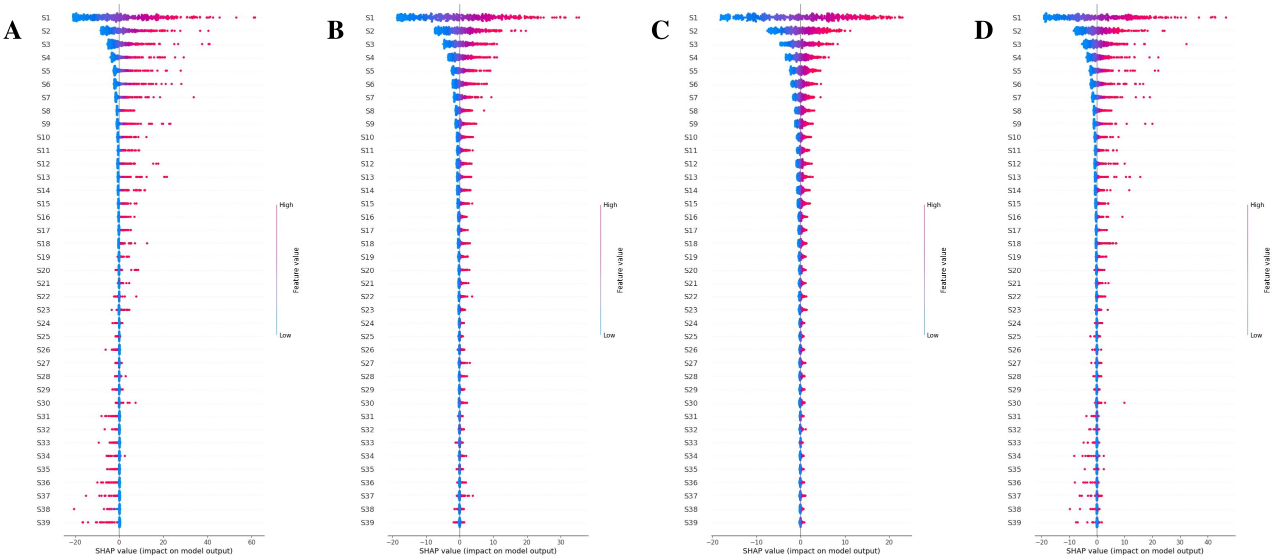


**Fig. 9. Feature importance adaptive neural network, trained with $\rho$ variable, via SHAP.** Feature importance of the adaptive neural network for $\rho = 0$ (A), $\rho = 35$ (B), $\rho = 1000$ (C) and $\rho$ variable (D). Here, the calculated shap values for A, B, C, and D are based on the adaptive neural network trained for variable recombination rates. The shap values were calculated for $500$ SFS. Each dot represents the shap value for a particular prediction of one feature, i.e. one SFS entry of one of the 500 SFS. The color indicates whether the particular SFS entry had a high or low value, and the position on the x-axis shows its significance, i.e., the extent to which it increased or decreased the $\theta$ estimate.

### Supplementary Note B: Neural Network Training Procedure

We considered two artificial neural networks: a linear dense feedforward neural network (no hidden layer) and a dense feedforward neural network with one hidden layer and an adaptive loss function. All neural networks take the site frequency spectrum $S$ as input and are trained for $\theta$ chosen uniformly in $(0,100)$. The neural networks for fixed recombination rates have been trained based on $2 \cdot 10^5$ simulations, for variable recombination rates with $6 \cdot 10^5$ simulations. For the linear neural network the division of simulations between training and validation was chosen to be 80% and 20%. Since the adaptive training procedure is similar to an ensemble training, in the sense that the process involves training multiple neural networks, a second validation set is required to evaluate in between the training steps. Therefore, an additional 20% is subtracted from the training set as a second validation set and the larger validation set is used to decide if another update of the adaptive loss function is needed. The test set is simulated separately and usually of size $4.9 \cdot 10^5$.

Hyperparameters are selected via a grid search. ReLU is used as activation function, the adam optimizer (28) with learning rate $0.001$, the number of hidden nodes is chosen as 200 and the batch size as $64$.

Preventing overfitting is always a challenging problem in ML. The risk of overfitting can be reduced not only in the training process by using methods such as regularization methods, but also via architectural design choices. First of all, the adaptive neural network has a lower model capacity compared to more sophisticated ML methods, making an overfit less likely. In addition, the adaptive training process is similar to ensemble learning with a termination condition (Eq 4) preventing the model from fitting the training data too well. Furthermore, each of the neural networks in the adaptive training procedure was trained using the regularization method *Early Stopping*. Training, validation and test errors were monitored. We observed no tendency of the model to overfit. For the generalization outside the training range, see also Figure S2.

### Supplementary Note C: Application with a Block Recombination Map

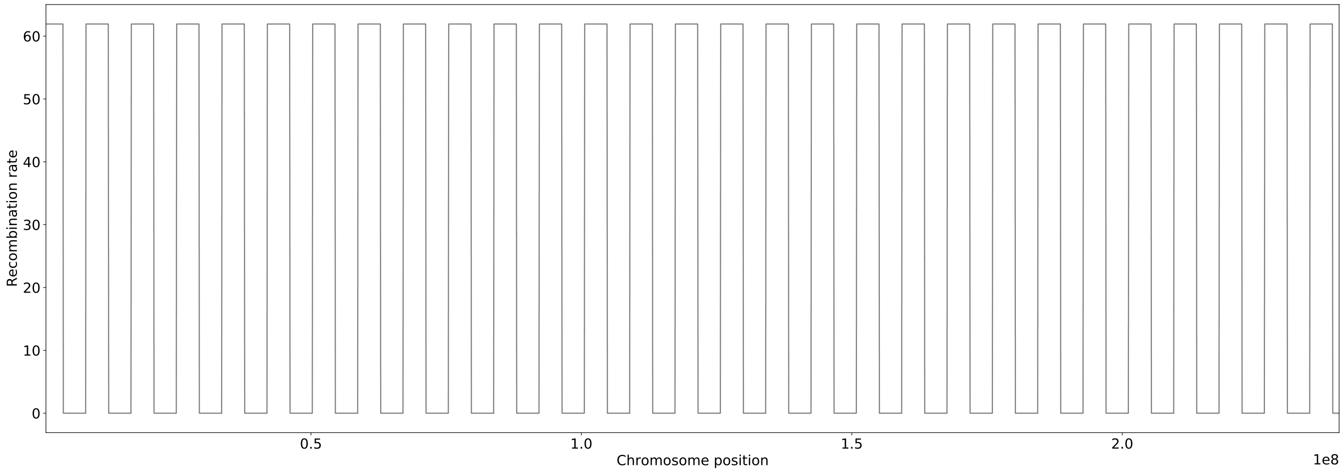


**Fig. 10. Block recombination map.** Recombination map with only two different recombination rates. The mean recombination rate matches that of chr2. Displayed recombination rates along the chromosome are based on a sliding window of $70$kb. This also ensures comparability with Fig 11.

To better illustrate the variability of the considered estimators along a chromosome we created an artificial recombination map with separated regions with low recombination. In this map the recombination rate jumps between $0$ and $61.9$ such that the mean recombination rate matches that of human chr2. This results in a *block recombination map* as displayed in Fig 10.

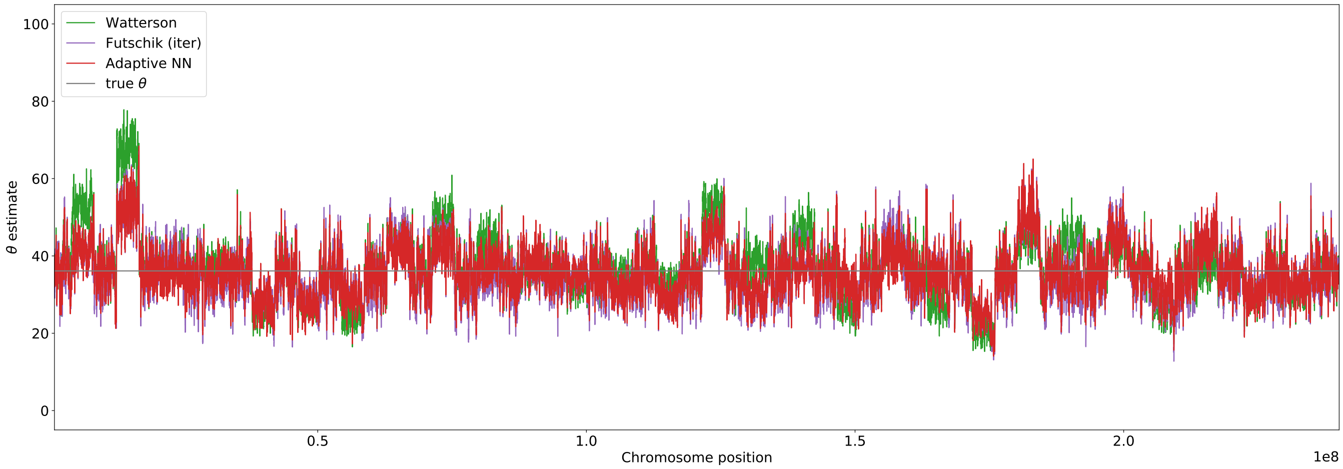


**Fig. 11. Estimation of $\theta$ for block recombination map.** SFS for $\theta$ estimation was calculated based on a sliding window of size $70$kb. Estimates for Watterson, iterative Futschik and the adaptive neural network are displayed. The neural network is trained with variable recombination rates. The underlying $\theta = 36.12$ is shown as the grey line.

Fig 11 displays the estimates of $\theta$ for Watterson's estimator, the iterated minimal MSE estimator (Futschik (iter)) and the adaptive neural network for the block recombination map (Fig 10). An extract not displaying the whole chromosome is shown in Fig 12 and for completeness the corresponding recombination map section in Fig 13. Figures 11 and 12 illustrate how the estimators adapt to the two recombination levels. Obviously, Watterson's estimator struggles for the parts without recombination. The iterative Futschik estimator is not particularly noticeable, in part because for high recombination rates the errors are generally much lower than for $\rho = 0$ and thus less detectable on the scale (see e.g. Fig 1). Of the estimators considered, the adaptive neural network uniform performs best.

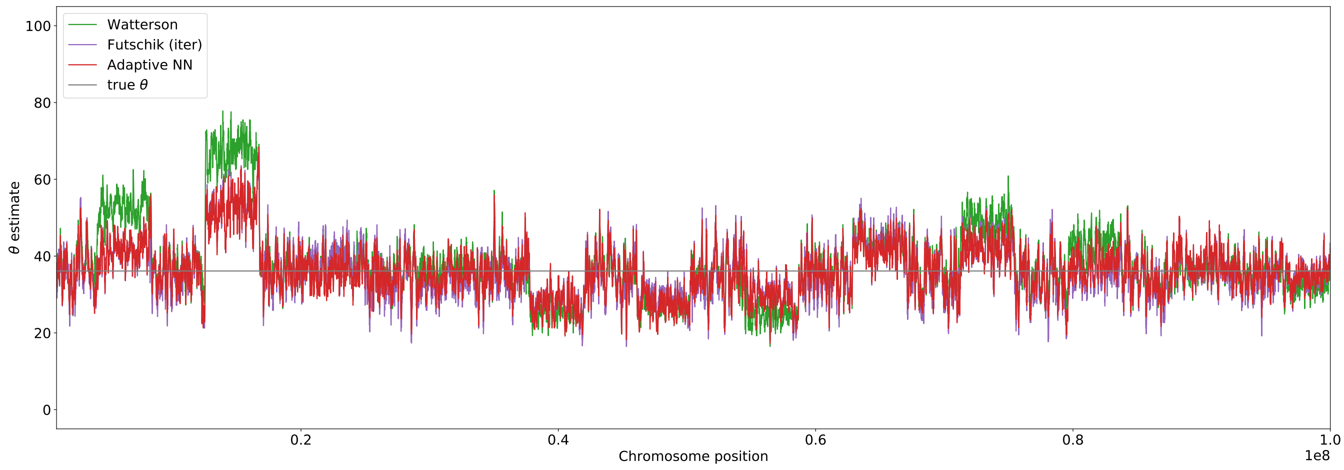


**Fig. 12. Extract of $\theta$ estimation for block recombination map.** SFS for $\theta$ estimation was calculated based on a sliding window of size $70$kb. Estimates for Watterson, iterative Futschik and the adaptive neural network are displayed. The neural network is trained with variable recombination rates. The underlying $\theta = 36.12$ is shown as the grey line.

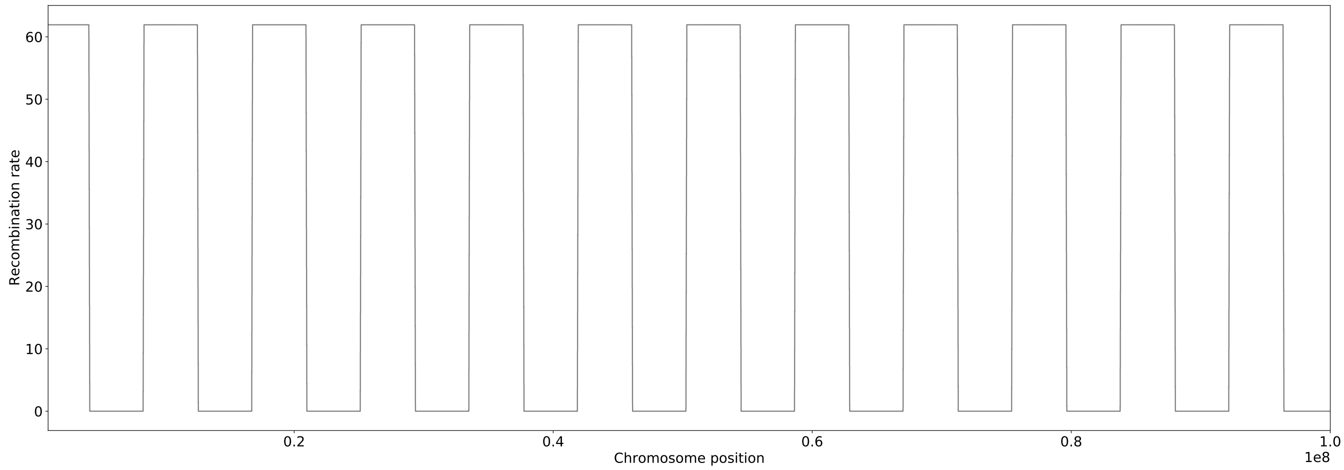


**Fig. 13. Extract of the block recombination map.** Extract of recombination map with only two different recombination rates. The mean recombination rate matches that of chr2. Displayed recombination rates along the chromosome are based on a sliding window of $70$kb. This also ensures comparability with Fig 12.

### S1 Supplemental Figure: Comparison of iterative and non-iterative Estimators for $\theta \in (0, 40]$

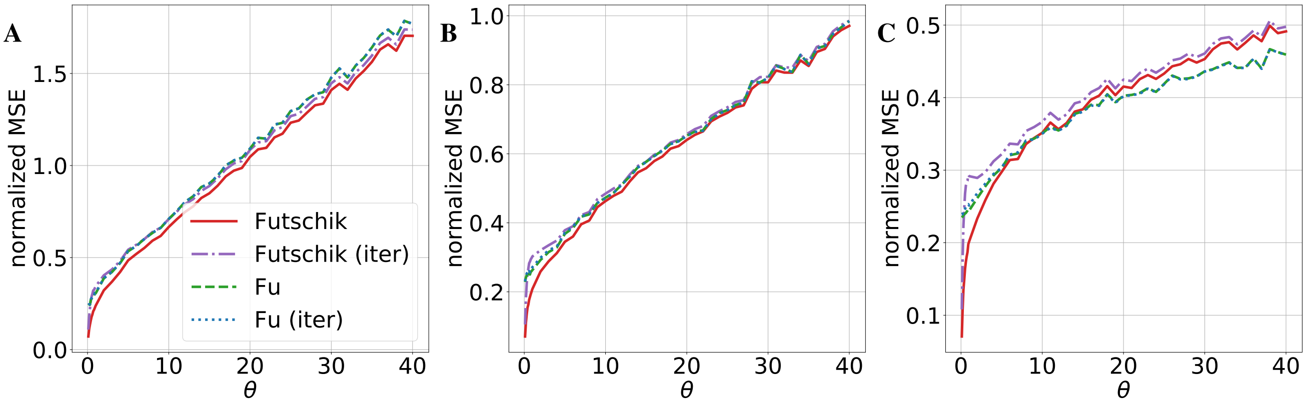


**Fig. S1. Iterative estimators vs. non-iterative estimators.** The normalized MSE for four different estimators are shown: The minimal MSE estimator (Futschik), the iterated minimal MSE estimator (Futschik (iter)), the minimal variance estimator (Fu) and the iterated minimal variance estimator (Fu (iter)). The same simulations as in Figure 1 have been used. A: recombination rate $\rho = 0$ B: $\rho = 35$ C: $\rho = 1000$

### S2 Supplemental Figure: Neural Network Performance outside the Training Range: $\theta$ in $(0, 500]$

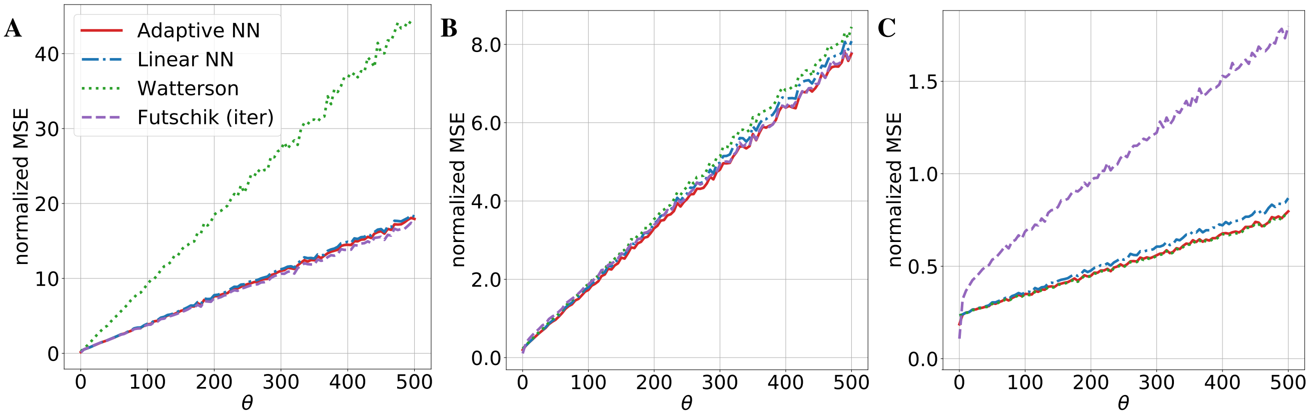


**Fig. S2. MSE for estimators up to $\theta = 500$.** The normalized MSE for four different estimators are shown: The iterated minimal MSE estimator (Futschik (iter)), Watterson's estimator, and two neural network estimators. A linear network without a hidden layer and an adaptive network with one hidden layer and 200 hidden nodes. All shown neural networks have been trained on data sets with the corresponding recombination rate $\rho$. For each subplot, we used msprime to generate a total of $18.9 \cdot 10^5$ independent simulations with sample size $n = 40$, recombination rate $\rho$, and 189 different mutation rates $\theta \in (0, 500]$. A: recombination rate $\rho = 0$ B: $\rho = 35$ C: $\rho = 1000$

### S3 Supplemental Figure: Normalized Bias of Estimators

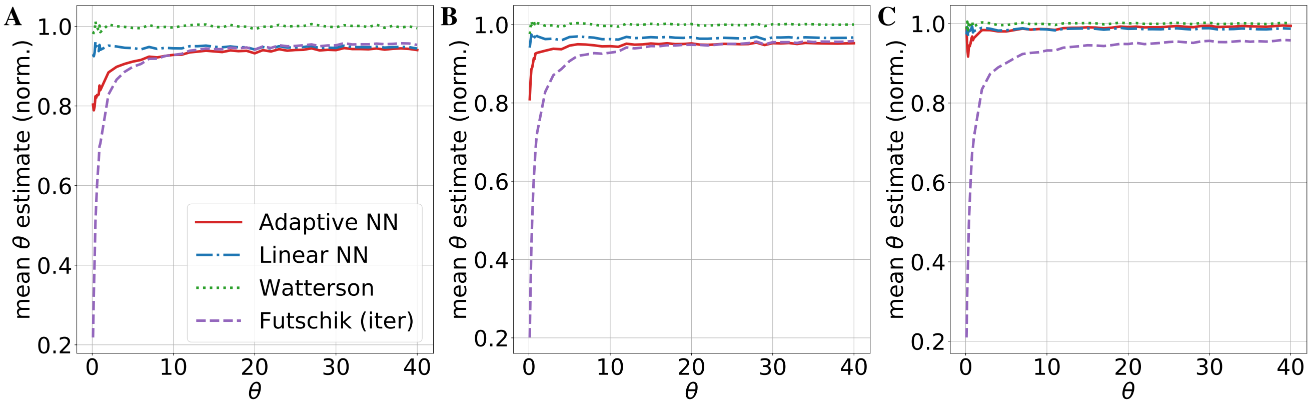


**Fig. S3. Normalized bias of estimators.** The normalized bias for four different estimators are shown: The iterated minimal MSE estimator (Futschik (iter)), Watterson's estimator, and two neural network estimators. A linear network without a hidden layer and an adaptive network with one hidden layer and 200 hidden nodes. All shown neural networks have been trained on data sets with the corresponding recombination rate $\rho$. The same simulations as in Figure 1 have been used. A: recombination rate $\rho = 0$ B: $\rho = 35$ C: $\rho = 1000$

### S4 Supplemental Figure: Coefficients of Fu's Estimator with Choice of Subsets for Adaptive Training

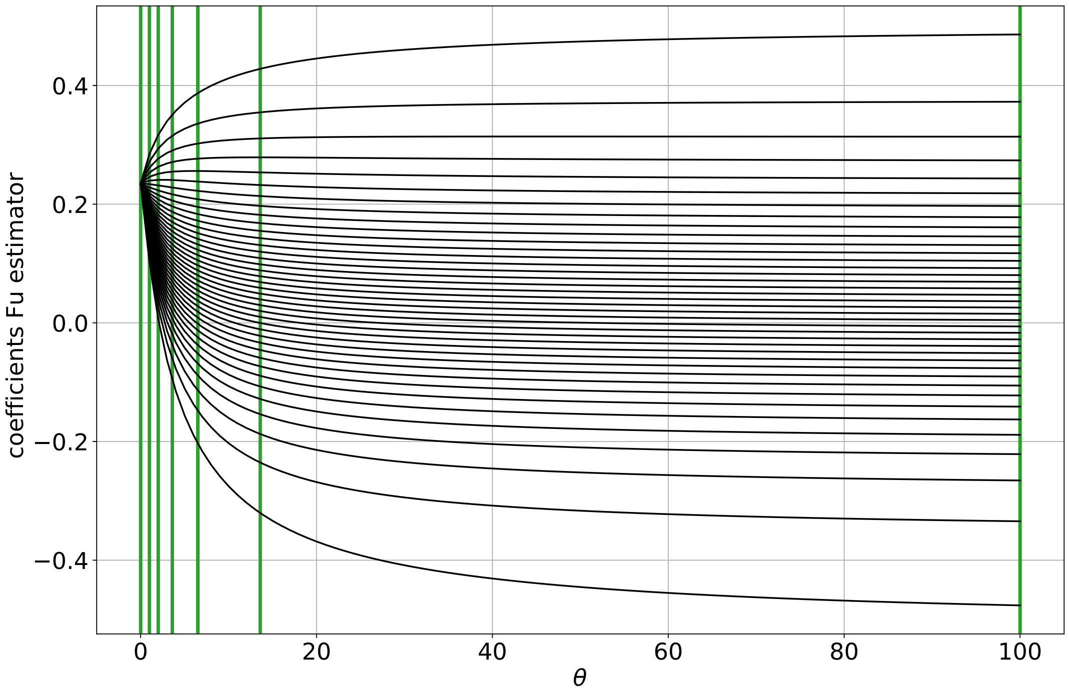


**Fig. S4. Coefficients of Fu's estimator with subsets for adaptive training procedure.** This figure shows the coefficients of non-interative Fu estimator for $n = 40$ (black) and the vertical lines represent the boundaries of the classes used in the adaptive training procedure for $n = 40$ (green).

### S5 Supplemental Figure: Visualization of the Adaptive Neural Network Architecture

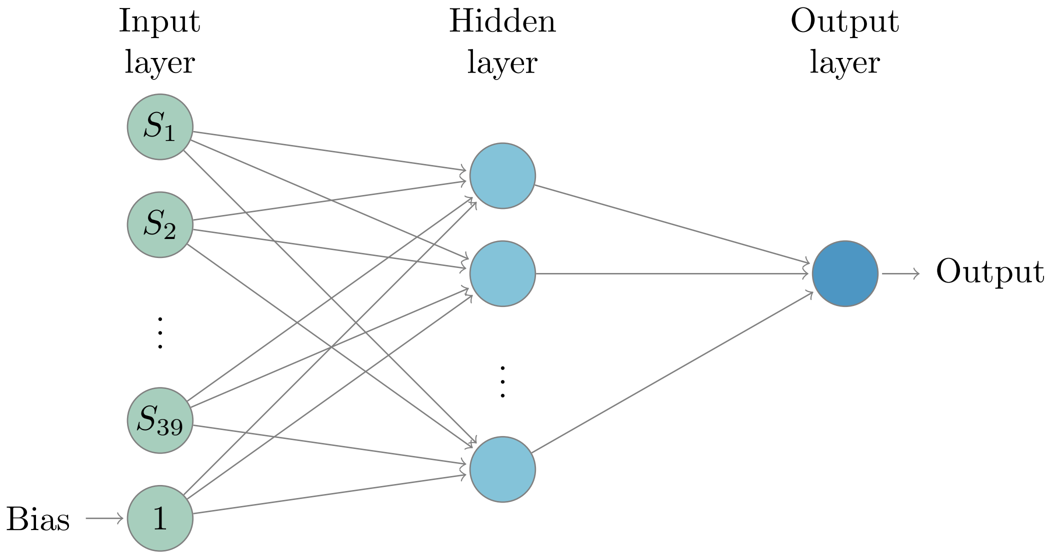


**Fig. S5. Adaptive neural network architecture.** The adaptive neural network considered in this publication is a dense feedforward neural network with one hidden layer, the site frequency spectrum as input and one bias node.

### S6 Supplemental Figure: Example for Mutations along a Coalescent Tree

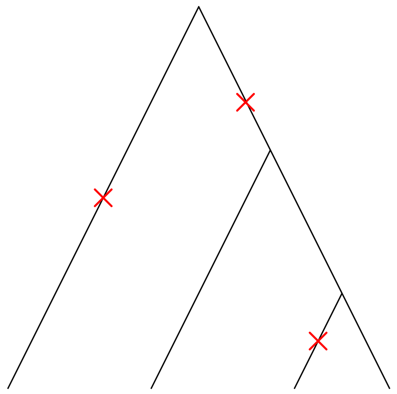

$$S = (S_1, S_2, S_3) = (2, 0, 1)$$

**Fig. S6. Coalescent tree with mutations.** Mutations along a coalescent tree for $n = 4$ and the corresponding site frequency spectrum $S$.

### S7 Supplemental Figure: Robustness of the Adaptive Neural Network and the CNN by Flagel et al.

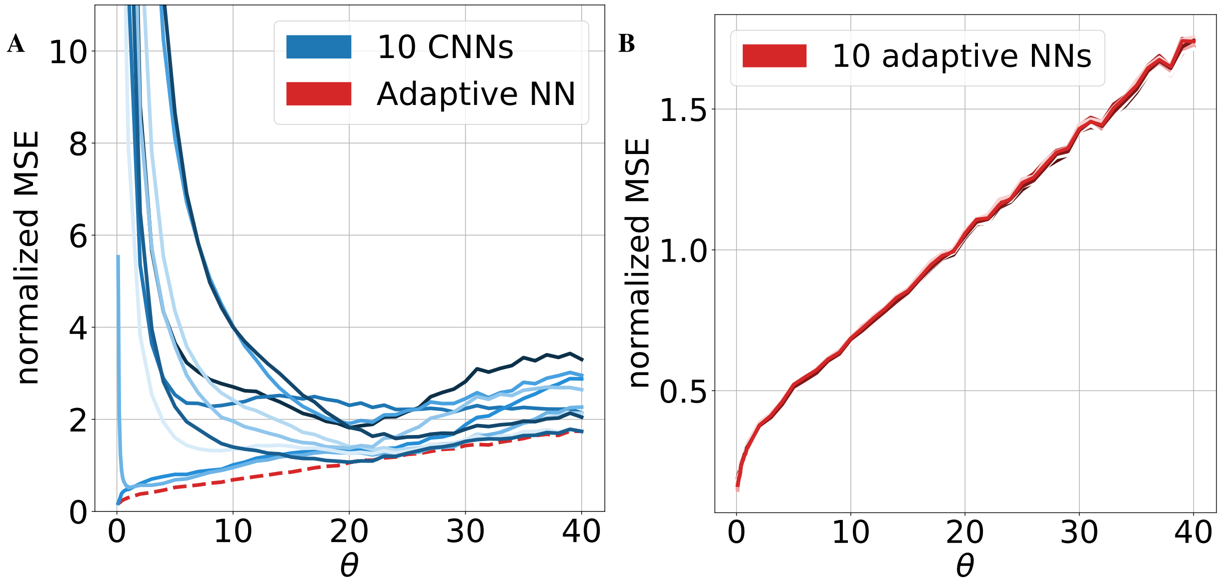


**Fig. S7. Robustness of neural networks.** Robustness for the CNN by Flagel et al. (21) (A) and the adaptive neural network (B). In each subplot, ten trained CNNs or adaptive neural networks, respectively, are displayed in the scenario where the recombination rate $\rho = 0$. One adaptive neural network (dashed line) has been added to (A) to facilitate the comparison.
